# Probing nanoscale diffusional heterogeneities in cellular membranes through multidimensional single-molecule and super-resolution microscopy

**DOI:** 10.1101/2020.08.14.232587

**Authors:** Rui Yan, Kun Chen, Ke Xu

## Abstract

Diffusion properties notably determine the behavior of biomembranes. Here we report the concurrent nanoscale fine-mapping of membrane topography, diffusivity, and packing order in live mammalian cells through a synergy of single-molecule and super-resolution methods. By identifying a bright, lipophilic fluorescence turn-on probe that enables sustained single-molecule imaging of cellular membranes under stroboscopic excitation, we accumulate the positions and transient displacements of >10^6^ probe molecules to achieve super-resolution topography and diffusivity mapping. We thus determine a trend that the membrane diffusivity drops with increased lipid packing order when comparing the endoplasmic reticulum (ER) membrane, plasma membrane, and nanodomains induced by cholera toxin B. Utilizing our nanoscale mapping capability, we further unveil reduced diffusivity in the ER membrane at ER-plasma membrane contact sites. By next integrating spectrally resolved single-molecule imaging, we show this localized diffusion slowdown is not due to altered lipid packing order, but may instead be attributed to local protein crowding. Our integrated multidimensional single-molecule approach thus unveils and differentiates between nanoscale diffusional heterogeneities of different origins in live-cell membranes.

## INTRODUCTION

Diffusion properties both determine the behavior of biomembranes and provide a valuable window into their dynamic structures.^1–5^ However, it remains difficult to obtain a continuous fine map of local diffusivity for cellular membranes. Typical diffusivity measurements based on fluorescence recovery after photobleaching (FRAP) and fluorescence correlation spectroscopy (FCS) afford limited spatial resolutions and/or mapping capabilities.^6,7^ Although single-particle tracking (SPT) offers nanoscale resolutions,^4,8,9^ historically it relies on sparse sampling so that only a few particles are tracked for each cell. The recent rise of single-molecule localization microscopy (SMLM) super-resolution methods^10–13^ has inspired new ways to achieve SPT at high densities. Photoactivation-based approaches^14–16^ allow temporal separation of single molecules within the diffraction limit, thus enabling the tracking of thousands of molecules over time. Methods based on *in situ* fluorescent probe exchange^17–19^ further overcome the limited photoactivation cycles of each probe. However, emphasis has been on obtaining trajectories for single protein molecules, which diffuse slow and often do not uniformly sample the membrane.

We recently reported single-molecule displacement/diffusivity mapping (SM*d*M), for superresolution mapping of the intracellular diffusion rates of unbound proteins.^20^ In contrast to SPT, in which each particle is tracked over many frames as it visits random locations, SM*d*M flips the question to evaluate, for each fixed location, how different (yet identical) single particles move locally by accumulating their transient displacements in short time windows. This location-centered strategy is powerful for spatial mapping. By eliminating the need to track long trajectories, SM*d*M is also ideal for systems in which fluorophores only emit transiently. PAINT (points accumulation for imaging in nanoscale topography)^13^ presents such a case for SMLM of the lipid membrane: As single probe molecules randomly enter the membrane phase from the aqueous medium, they turn on fluorescence emission and reside for ~10 ms before exiting,^21^ thus allowing unbiased, high-density membrane sampling.^13^

Here we exploit the transient intra-membrane diffusion of single probe molecules in PAINT to achieve SM*d*M diffusivity mapping for cellular membranes. By identifying a bright fluorescence turn-on probe that enables sustained PAINT-type single-molecule imaging for >10^5^ frames under stroboscopic illumination, we accumulate the super-resolved positions and transient displacements of >10^6^ molecules in live-cell membranes. We thus achieve continuous fine mapping of the cellular membrane diffusivity to unveil heterogeneities at the nanoscale, and further integrate spectrally resolved SMLM (SR-SMLM)^22,23^ to examine the possible roles of lipid packing order and protein crowding in local diffusivity differences.

## RESULTS

For effective probing of both the topography and diffusion properties of cellular membranes, we started by identifying the commercial dye BDP-TMR-alkyne (Figure 1a) as an optimal probe. BDP-TMR has been reported as a water-soluble dye with a high affinity to lipid bilayers.^24^ We found BDP-TMR-alkyne exhibited ~4-fold fluorescence turn-on in organic phases vs. in the aqueous medium (Figure 1b). When added into the cell medium at 3.3 nM, it emitted strongly over low backgrounds as single molecules transiently entered cellular membranes (Figure 1c). In comparison to Nile Red, the typical PAINT probe for lipid membranes,^13,25,26^ BDP-TMR-alkyne was notably brighter and yielded superior single-molecule images (Figure 1c-e). When excited by stroboscopic pulses of *τ* = 1 ms duration to minimize motion blur, a substantial fraction of the molecules emitted ~1,000 photons (Figure 1e), thus enabling reliable three-dimensional (3D) localization through single-molecule image shape.^27^ Similar to Nile Red, BDP-TMR-alkyne dynamically partitioned between the medium and membrane phases so that single molecules stayed in the membrane for ~10 ms^21^, a timescale ideal for the <10 ms displacement time window of SM*d*M (below). The rapid membrane-medium exchange created a steady state, so that continuous imaging was readily achieved for >1.6×10^5^ frames with no appreciable drop in the singlemolecule count (Figure 1f), from which 10^6^-10^7^ molecules were accumulated across the view. It is further worth mentioning that these results were achieved in the regular cell media without the need for an oxygen scavenger, other additives, or photoactivation.

**Figure 1.**
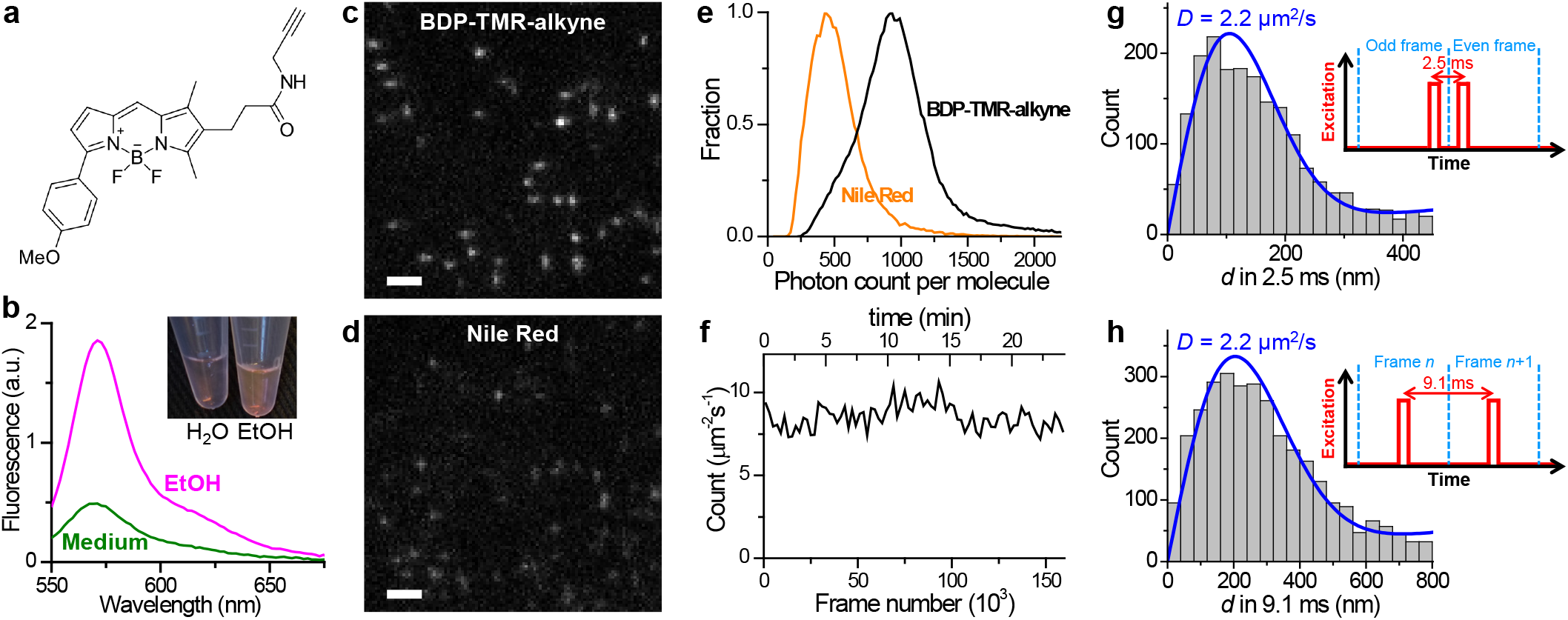
BDP-TMR-alkyne for stroboscopic PAINT and SM*d*M of cellular membranes. (a) Chemical structure of BDP-TMR-alkyne. (b) Fluorescence emission spectra of 3.3 nM BDP-TMR-alkyne in the aqueous cell medium (green curve) vs. in ethanol (magenta curve). Inset: photo of 10 μM solutions of BDP-TMR-alkyne in water vs. in ethanol. (c,d) Typical single-molecule images of BDP-TMR-alkyne (c) vs. Nile Red (d) in cell membranes recorded under identical experimental conditions with stroboscopic excitation pulses of *τ* = 1 ms duration. A cylindrical lens was applied to introduce image elongations in vertical or horizontal directions for molecules of different depths. (e) Distribution of single-molecule photon counts for the two cases. (f) Time-dependent count of the detected BDP-TMR-alkyne single molecules per μm^2^ per second at the plasma membrane of a COS-7 cell, showing no noticeable drop over 1.6×10^5^ frames (24 min at 110 fps). (g,h) Typical distributions of transient single-molecule displacement *d* of BDP-TMR-alkyne at the COS-7 cell plasma membrane under two stroboscopic excitation schemes (insets): repeated tandem excitation pulses at a Δ*t* = 2.5 ms center-to-center separation (g), vs. excitation at the middle of every frame at 110 fps (Δ*t* = 9.1 ms) (h). Blue curves: MLE (maximum likelihood estimation) fits to our model, with resultant diffusion coefficient *D* labeled. Scale bars: 2 μm (c,d).

For SM*d*M, we implemented two different excitation schemes while continuously recording in the wide-field at 110 frames per second (fps). Whereas the application of asymmetrically timed pulse pairs across tandem frames^20^ at a Δ*t* = 2.5 ms center-to-center separation helped limit the transient singlemolecule displacement *d* to <~200 nm (Figure 1g) for more localized statistics, exciting at the middle of each frame (Figure 1h) allowed *d* in Δ*t* = 9.1 ms time windows to be extracted between consecutive frames. Fitting the *d* distributions obtained under the two excitation schemes to a modified two-dimensional (2D) random-walk model^20^ gave similar typical diffusion coefficients of ~2.2 μm^2^/s for cell plasma membranes (Figure 1gh), comparable to that previously measured by SPT for other lipophilic dyes.^15,28^ We further obtained comparable results for excitation pulses of different duration *τ* (Figures S1 and S2), thus demonstrating the robustness of our measurements with varied parameters.

The possibility to accumulate a large amount of single-molecule data through stroboscopic PAINT of BODIPY-TMR-alkyne enabled continuous fine-mapping of membrane topography and diffusivity. Resultant 3D-SMLM images (Figure 2ab, Figure S1, and Figure S2) well resolved how the dorsal (top) plasma membrane raised out of the focal range when far away from the cell edge, as well as how the intracellular organelle membranes, *e.g*., that of the endoplasmic reticulum (ER), undulated in height and possibly contacted the plasma membrane (arrowheads in Figure 2b). The ventral (bottom) plasma membrane was lowly labeled owing to its inaccessibility to the probe-containing medium, similar to our observations for other PAINT probes.^26,29^ Meanwhile, spatially binning the accumulated transient single-molecule displacement *d* into 100×100 nm^2^ bins enabled their individual fitting^20^ to obtain the local diffusivity *D*, hence color-coded SM*d*M super-resolution maps (Figure 2c, Figure S1, and Figure S2). Relatively uniform *D* with typical values of 2.0-2.5 μm^2^/s was found for the plasma membrane. Treating the cell with cholera toxin B subunit (CTB), a cross-linker of the ganglioside GM1, gave rise to ~200 nm-sized nanodomains of substantially reduced local diffusivity down to ~1 μm^2^/s (Figure 2e), which partially co-localized with CTB (arrows in Figure 2de). This result echoes our recent SR-SMLM results with Nile Red, which showed that CTB induces similarly-sized nanodomains of low chemical polarity in the plasma membrane, attributable to increased lipid packing order due to the local sequestration of GM1 and cholesterol.^23,26^ Cholesterol-induced packing order and reduced membrane fluidity are well documented at the bulk level for both cell and model membranes.^30–33^

**Figure 2.**
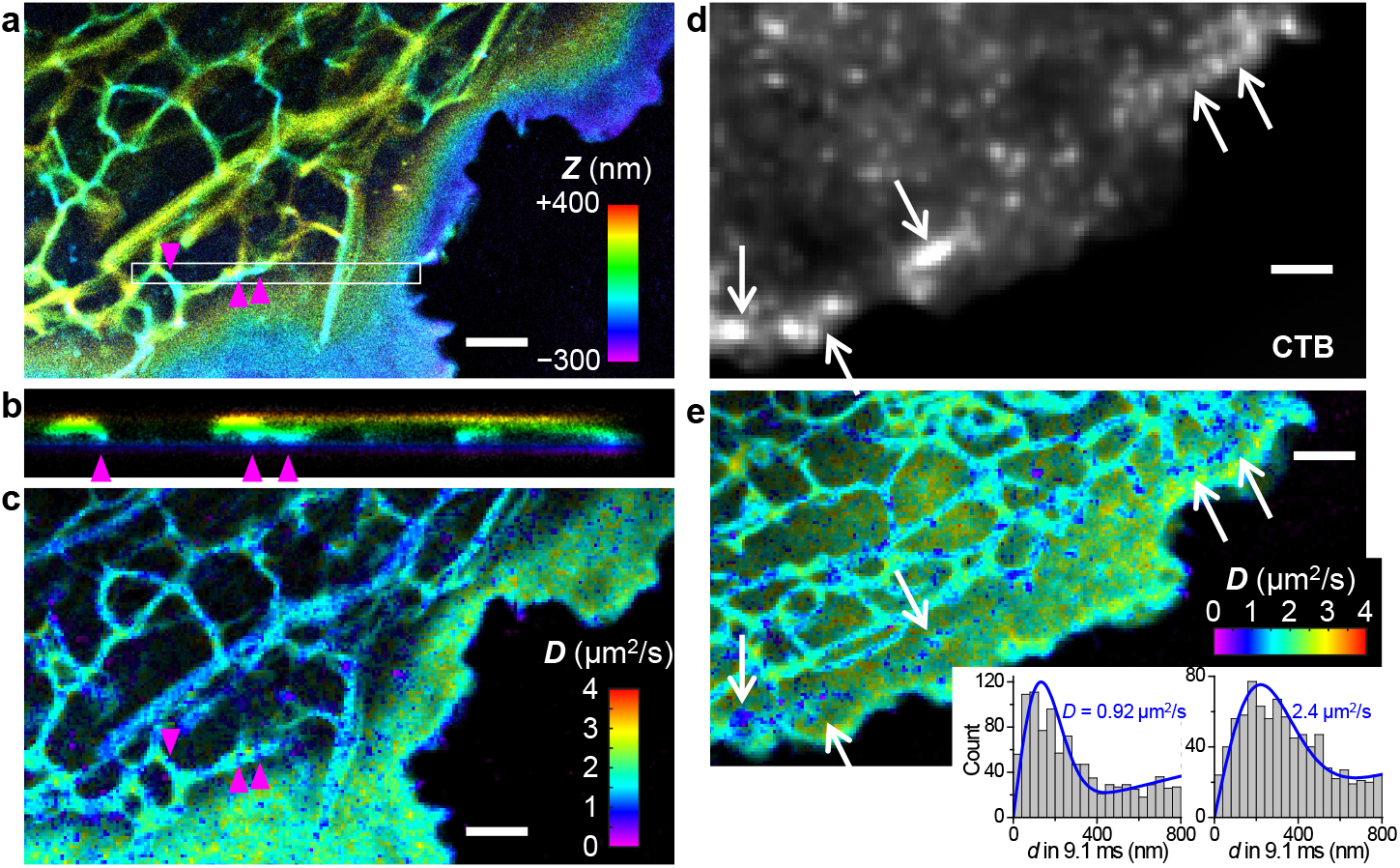
Concurrent 3D-SMLM and SM*d*M diffusion mapping of cellular membranes through stroboscopic PAINT of BDP-TMR-alkyne. (a) 3D-SMLM image obtained in a live COS-7 cell. Color presents depth (*z*); purple/blue: closest to the coverslip; red/yellow: farthest away. (b) Cross-sectional view along the box in (a) in the *xz* direction. Arrowheads point to possible contacts between ER and the ventral plasma membrane. (c) SM*d*M diffusivity map constructed from the same single-molecule data. Color presents local diffusivity *D*. (d) Epifluorescence image of dye-tagged CTB applied to another COS-7 cell. (e) BDP-TMR-alkyne SM*d*M diffusivity map of the same cell. Insets: typical distributions of single-molecule displacements for the low-diffusivity nanodomains (left) versus other parts of the plasma membrane (right), and their MLE fits for *D* (blue curves). For both cells, stroboscopic excitation of *τ* = 1 ms duration was applied at the middle of every frame at 110 fps (Δ*t* = 9.1 ms). Scale bars: 2 μm.

For the ER tubule membrane, typical *D* values appeared slightly lower than the plasma membrane (Figure 2ce), apparently against trend with their lower cholesterol level and packing order.^26,34,35^ 2D plotting of single-molecule displacements showed a preferred angular distribution along the tubule (Figure 3a), indicating the isotropic 2D fitting model we originally used unsuitable. This observation underscores a geometrical effect that underestimates motion perpendicular to the tubule length: in the extreme case, a molecule that traveled a round trip along the tubule circumference in Δ*t* would give a zero displacement.

**Figure 3.**
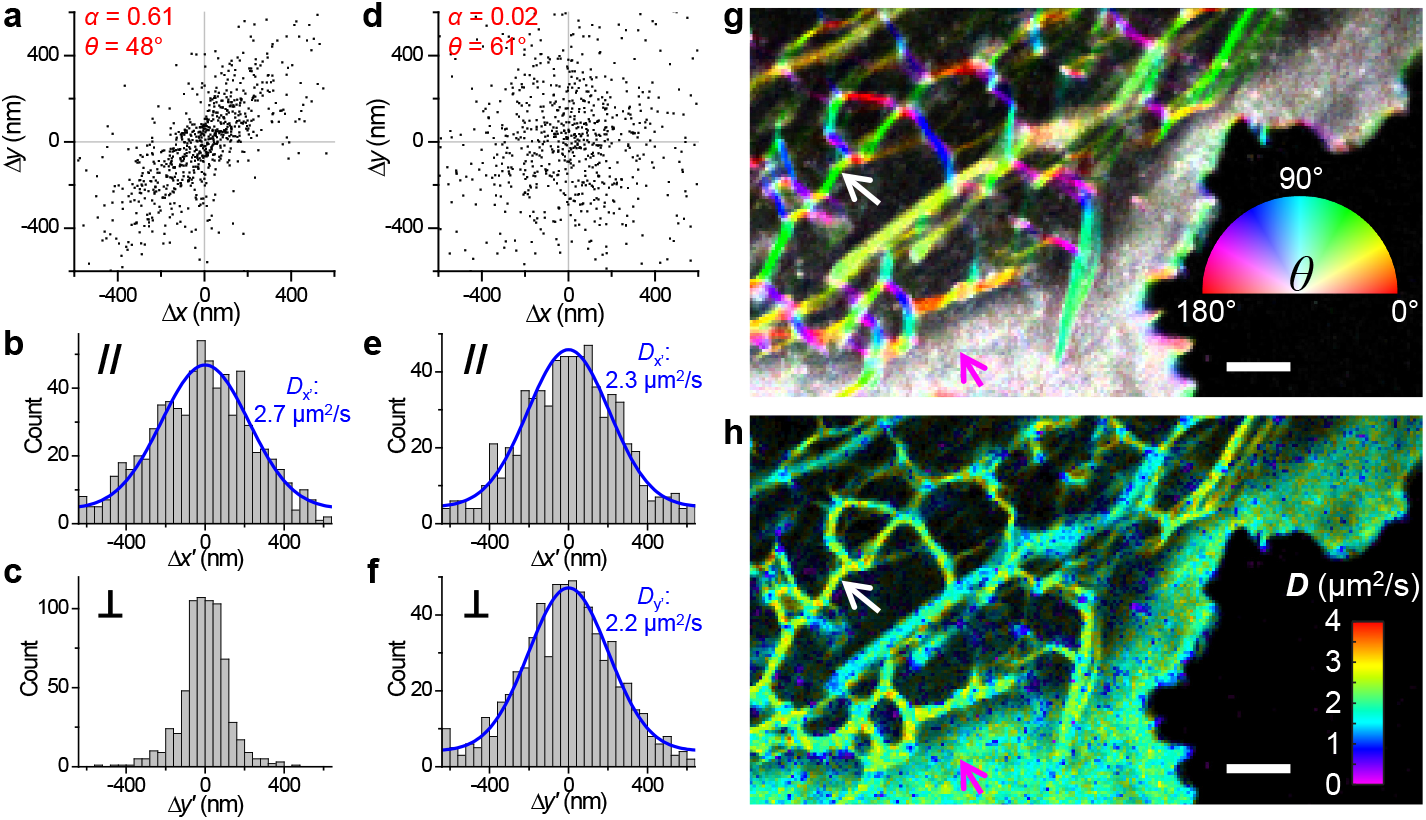
Principal-direction SM*d*M (pSM*d*M) analysis. (a) 2D plot of single-molecule displacements in Δ*t* = 9.1 ms for BDP-TMR-alkyne at an ER tubule [white arrows in (g,h)]. A principal direction *θ* of 48° is calculated with an anisotropy *α* of 0.61. (b,c) 1D distributions of the displacements in (a) projected along (b) and perpendicular (c) to *θ*. Blue curve in (b): MLE fit to an 1D diffusion model with resultant *D* = 2.7 μm^2^/s. (d) 2D plot of singlemolecule displacements in Δ*t* = 9.1 ms for BDP-TMR-alkyne at the plasma membrane [magenta arrows in (g,h)]. *α* = 0.02 and *θ* = 61°. (e,f) 1D distributions of the displacements in (d) projected along (e) and perpendicular (f) to *θ*. Blue curves: MLE fits to a 1D diffusion model, with resultant *D* = 2.3 and 2.2 μm^2^/s. (g) Color map presenting the calculated local *θ* (hue; inset color wheel) and α (color saturation; range 0-0.5) for the same region as Figure 2c. (h) pSM*d*M diffusivity map based on 1D diffusion fits along local principal directions. Scale bars: 2 μm (g,h).

We reason that the true diffusivity may still be obtained by examining displacements *along* the tubule. To this end, we started by determining the local displacement anisotropy *α* and principal direction *θ* (Figure 3a; Methods). Projecting single-molecule displacements along and perpendicular to *θ* gave contrasting distributions: the former (Figure 3b) represents nonrestraint diffusion along the tubule, which could be fit to a modified 1D diffusion model^20^ to extract a *D* value (Figure 3b), whereas the latter (Figure 3c) is convolved with tubule width. In comparison, displacements at the plasma membrane were isotropic (Figure 3d), and comparable *D* values were obtained when the 1D diffusion model was applied to projections along and perpendicular to the near-arbitrarily defined *θ* (Figure 3ef). Applying the above analysis to every spatial bin showed well-defined directional preferences along ER tubules in the resultant *θ*-*α* color maps (Figure 3g and Figure S1). Projecting the single-molecule displacements in each bin along its *θ* direction next enabled their individual fitting to the 1D diffusion model. Resultant principal-direction SM*d*M (pSM*d*M) diffusivity maps (Figure 3h, Figure S1, and Figure S2) showed *D* values higher in the ER membrane (~2.7 μm^2^/s) than in the plasma membrane (~2.2 μm^2^/s), in line with their expected lower lipid packing order and hence possibly higher fluidity. This strategy, however, did not overcome the complex, cristae geometry of mitochondrial membranes, for which *D* values were still underestimated (*e.g*., marked by asterisks in Figure 4), awaiting future investigations.

**Figure 4.**
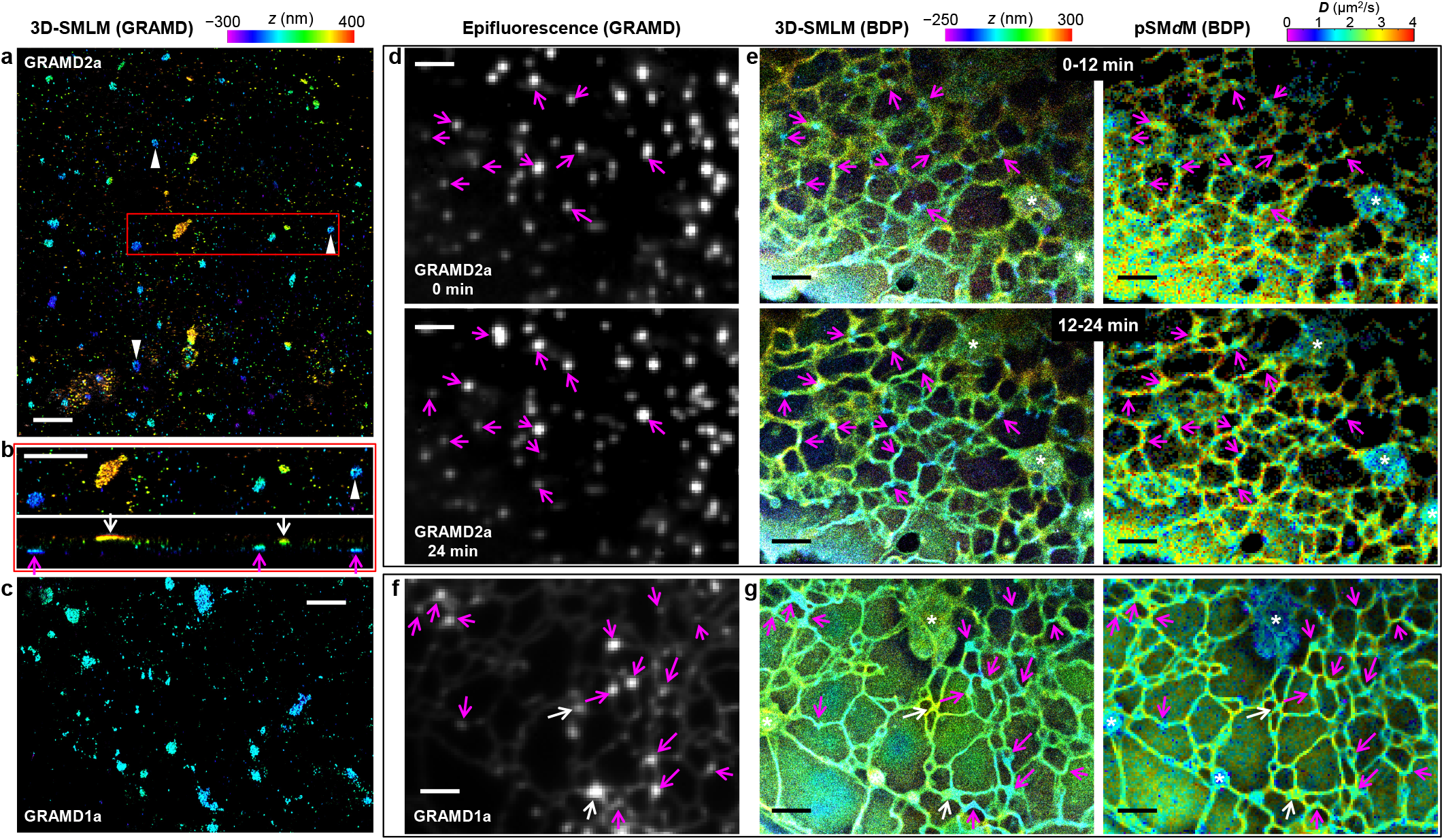
Concurrent SM*d*M and 3D-SMLM, together with fluorescent protein markers, indicate reduced membrane diffusivity at ER-PM contact sites. (a) 3D-SMLM image of immunolabeled GRAMD1a-GFP expressed in a COS-7 cell. Color presents depth (color scale on the top). (b) Zoom-in of the red box in (a) (top) and its vertical view in the *xz* direction (bottom). Magenta and white arrows point to apparent contact sites at the ventral and dorsal plasma membranes, respectively. (c) 3D-SMLM image of immunolabeled GRAMD1a-GFP at the ventral plasma membrane of another COS-7 cell. (d) Live-cell epifluorescence images of GRAMD2a-GFP expressed in a COS-7 cell, taken before (top) and after (bottom) stroboscopic PAINT of BDP-TMR-alkyne. (e) 3D-SMLM (left) and pSM*d*M (right) images (separate color scales on the top) of the same region via stroboscopic PAINT of BDP-TMR-alkyne, for two consecutive 12-min periods. Data were acquired with tandem excitation pulses of *τ* = 1 ms duration at Δ*t* = 2.5 ms. Arrows in (d,e) point to example GRAMD2a-positive ER-PM contacts. Asterisks mark mitochondria, for which *D* values were underestimated due to complex membrane geometries. (f) Epifluorescence image of GRAMD1a-GFP expressed in another live COS-7 cell. (g) 3D-SMLM (left) and pSM*d*M (right) images of BDP-TMR-alkyne of the same region. Data were acquired with excitation pulses of *τ* = 1 ms duration at the middle of each frame at Δ*t* = 9.1 ms. Magenta and white arrows in (f,g) point to example GRAMD1a-positive contact sites at the ventral and dorsal membranes, respectively. Scale bars: 2 μm.

Closer examination of both the isotropic and principal-direction SM*d*M data showed the ER diffusivity sporadically exhibit local reductions in *D*, often at tubule junctions and termini close to the plasma membrane (Figures 2, 3, S1, and S2). To examine the possibility that these low-diffusivity foci corresponded to ER-plasma membrane (ER-PM) contact sites,^36–38^ we expressed in the cells green fluorescent protein (GFP)-tagged GRAMD1a or GRAMD2a, which localizes to a subset of the EP-PM contacts.^39^ 3D-SMLM through immunolabeling GFP in fixed cells showed GRAMD2a as ~100–500 nm patches at both the dorsal and ventral plasma membranes (Figure 4ab), often of slightly elongated shapes and occasionally with hollow centers (arrowheads). GRAMD1a similarly appeared as nanoscale patches, but with somewhat higher structural heterogeneities (Figure 4c).

To co-visualize these EP-PM contact site markers with membrane topography and diffusivity in live cells, we recorded epifluorescence images of GRAMD2a-GFP both before and after a long stroboscopic PAINT recording of BDP-TMR-alkyne over 24 min (Figure 4d). The single-molecule data were divided into two image stacks for the 0-12 min and 12-24 min periods, and each processed into matching 3D-SMLM and pSM*d*M images (Figure 4e). We thus found the GRAMD2a-GFP foci before and after the single-molecule recording separately matched the first and second 3D-SMLM images for ER tubule junctions/termini of low *z* values (arrows), indicative of contacts to the ventral plasma membrane. Notably, the concurrently obtained SM*d*M images showed substantially (~30%) reduced *D* at the same sites (Figure 4e), thus indicating that the ER-PM contacts maintained locally reduced membrane diffusivity as they reorganized in the live cell. In a different experiment on a thinner cell, GRAMD1a-GFP labeling (Figure 4f) and 3D-SMLM (Figure 4g) helped identify ER-PM contact sites at both the ventral (magenta arrows) and dorsal (white arrows) plasma membranes, and SM*d*M indicated similar diffusion slowdowns for both cases (Figure 4g).

To understand whether the observed diffusion slowdown at ER-PM contact sites could be related to changes in the lipid packing order as we showed above for the CTB treatment, we next integrated SM*d*M with SR-SMLM^22,23^ using the solvatochromic membrane probe Nile Red, which detects variations in lipid packing order through shifts in emission spectrum due to changes in the membrane chemical polarity.^23,26,35^ Although the lower single-molecule brightness of Nile Red (Figure 1de) was not optimal for 3D localization, it worked well for SM*d*M with stroboscopic excitation pulses of *τ* = 2 ms duration (Figure 5a). Reduced local diffusivity was thus similarly observed at ER-PM contact sites marked by GRAMD2a-GFP (Figure 5c). Meanwhile, the collected single-molecule emission spectra at these sites were identical to adjacent ER regions (Figure 5bd), indicating no significant changes in the local lipid order.^23,26^ In comparison, markedly bluer emission was observed for the plasma membrane (Figure 5bd) accompanying its lower diffusion rate, as expected from cholesterol-associated packing order.^26,35^ The ER-PM contact sites are characterized by a wealth of trans-ER-membrane proteins that interact with plasma-membrane components to achieve key functions as lipid exchange and Ca^2+^ regulation,^36–38^ and electron microscopy indicates high protein densities.^40,41^ It is thus possible that macromolecular crowding in the ER membrane could create a maze that impedes diffusion^1,42–44^ without altering the local lipid packing order.

**Figure 5.**
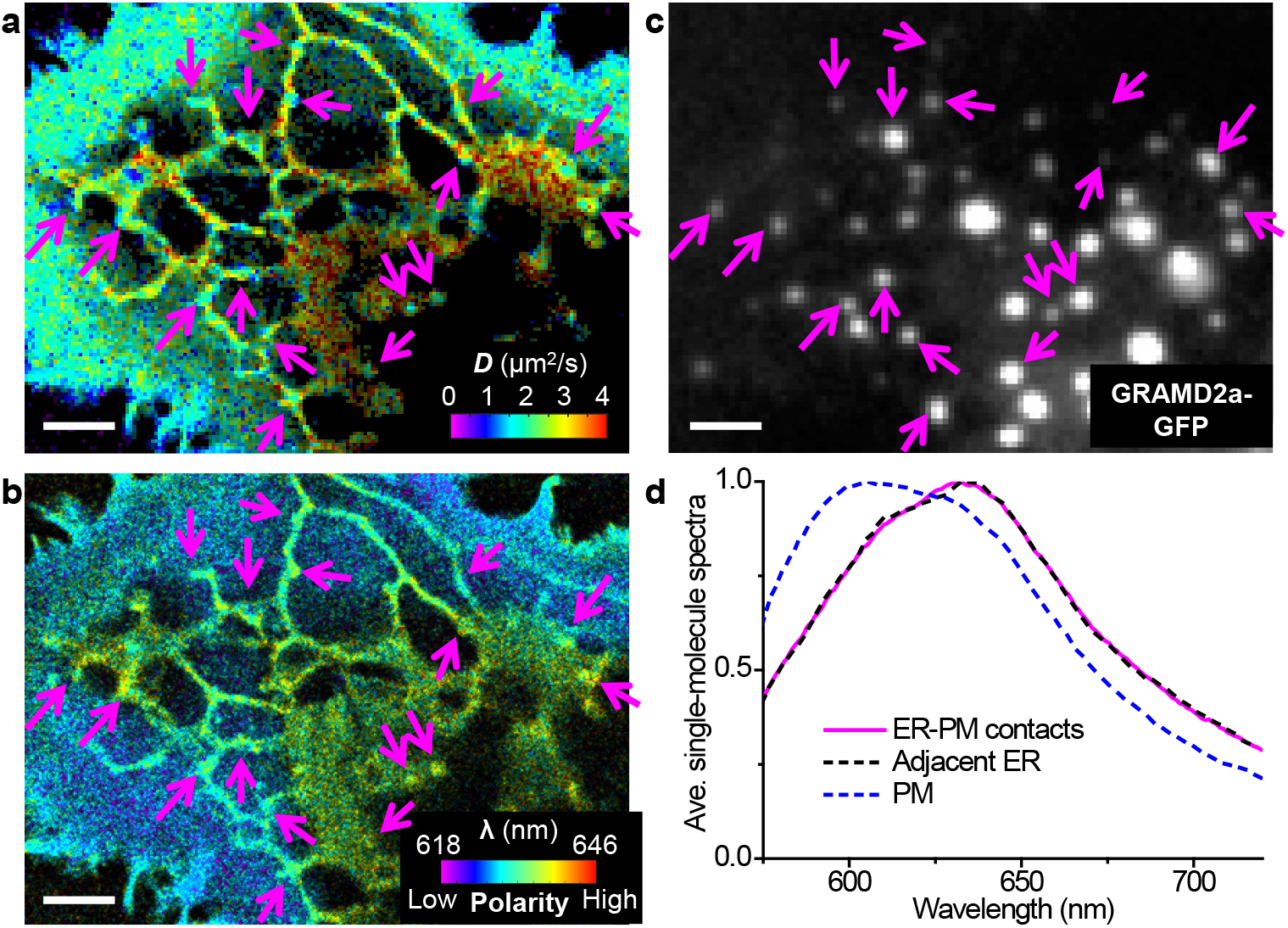
Concurrent SM*d*M and SR-SMLM of Nile Red indicate that the reduced diffusivity at ER-PM contacts is not associated with significant changes in lipid order. (a,b) Concurrently acquired pSM*d*M (a; color presents local diffusivity) and SR-SMLM (b; color presents the spectral mean of locally accumulated single molecules) images of Nile Red in a live COS-7 cell. Data were acquired with tandem excitation pulses of *τ* = 2 ms duration at Δ*t* = 2.5 ms. (c) Epifluorescence image of GRAMD2a-GFP expressed in this cell. Arrows point to example GRAMD2a-positive ER-PM contact sites where substantial reductions in *D* are noticed in the pSM*d*M image. (d) Averaged local singlemolecule spectra at the ER-PM contacts (magenta line) vs. adjacent ER regions (black dashed line), as well as at the plasma membrane (blue dashed line). Scale bars: 2 μm (a-c).

## DISCUSSION

By identifying a bright, lipophilic fluorescence turn-on probe that enabled sustained stroboscopic PAINT experiments over long periods, we accumulated high densities of single-molecule positions and displacements in cellular membranes to achieve nanoscale fine-mapping of membrane topography and diffusivity. Analysis of the local principal displacement direction helped overcome a geometrical effect that underestimated the diffusion in ER tubules; together with CTB results, we thus determined a trend that the membrane diffusivity drops with increased local lipid packing order. Utilizing the fine-mapping capability of SM*d*M while further incorporating 3D localization and specific fluorescent protein markers, we next unveiled reduced diffusivity in the ER membrane at ER-PM contact sites. By next integrating SR-SMLM, we showed that this effect was not due to altered local lipid order, hence implicating protein crowding as a possible source of the diffusion slowdown. Experimental work on cells and model lipid bilayers, together with theory, has well addressed how protein crowding impedes membrane diffusion at the bulk level.^1,42–44^ However, as previous diffusion measurements offer limited mapping capabilities, how such effects might *locally* modulate membrane diffusivity remains elusive. By offering fine maps of local diffusivity, SM*d*M unveiled local diffusion slowdowns at the ER-PM contact sites, consistent with their high protein densities observed in electron microscopy.^40,41^ Such effects may help stabilize the contact sites and assist material exchanges^36–38^ between the ER and the plasma membrane, the potential consequences of which call for future experimental and theoretical investigations. Together, as a timely integration of emerging modes of functional single-molecule and super-resolution approaches,^45^ our work thus unveiled and differentiated between nanoscale diffusional heterogeneities of different origins in live-cell membranes.

## ACKNOWLEDGMENTS

We thank W. Li and S. Moon for discussion. This work was supported by the National Science Foundation (CHE-1554717), the National Institute of General Medical Sciences of the National Institutes of Health (DP2GM132681), the Beckman Young Investigator Program, and the Packard Fellowships for Science and Engineering. K.X. is a Chan Zuckerberg Biohub investigator.

