## Supporting Information for "Probing nanoscale diffusional heterogeneities in cellular membranes through multidimensional single-molecule and super-resolution microscopy"

### Materials and Methods

**Cell culture.** COS-7 cells (University of California Berkeley Cell Culture Facility) were maintained in Dulbecco's Modified Eagle Medium supplemented with 10% fetal bovine serum and 1% non-essential amino acids. Two days prior to imaging, cells were plated onto 18-mm diameter glass coverslips that were pretreated with hot piranha solutions ($H_2SO_4$:$H_2O_2$ at 3:1). Transfection of GRAMD1a-AcGFP and GRAMD2a-AcGFP [kind gifts from Prof. Jodi Nunnari (Besprozvannaya *et al.* 2018)] was performed one day after plating. 300-500 ng plasmid was used per sample with the standard Lipofectamine 3000 protocol (Thermo Fisher). Before imaging, the coverslip was transferred to a holder (CSC-18, Bioscience Tools) compatible with the microscope stage. Imaging medium was Leibovitz's L-15 or a MOPS-based buffer (Hibernate A Low Fluorescence, BrainBits), with similar results observed. BDP-TMR-alkyne (Lumiprobe) or Nile Red (Acros Organics) were prepared as 1 mM stock solutions in DMSO, and diluted into the imaging medium to a final concentration of 3.3 nM. The dye-added imaging medium was added to the sample and remained unchanged throughout imaging. For CTB treatment, cells were incubated with 1 μg/mL Alexa Fluor 488-conjugated CTB (C34775, Invitrogen) in the culture medium for ~5 min, and then washed twice with the imaging medium before imaging.

**Concurrent SM*d*M and 3D-SMLM.** Concurrent SM*d*M and 3D-SMLM of BDP-TMR-alkyne was achieved *via* modifications to a homebuilt system (Wojcik *et al.* 2015) based on a Nikon Eclipse Ti-E inverted optical microscope. Lasers at 488, 560, and 647 nm were independently modulated by an acousto-optic tunable filter (97-03151-01, Gooch & Housego) that was driven by an 8-channel RF synthesizer (97-03926-12, Gooch & Housego). The modulated laser beams were focused onto the back focal plane of an oil-immersion objective lens (Nikon CFI Plan Apochromat λ 100x, numerical aperture: 1.45) toward the edge, thus entering the sample slightly below the critical angle to illuminate a ~1 μm depth. For 3D localization, a cylindrical lens was used to induce elongations of single-molecule images in the vertical and horizontal directions for molecules below and above the focal plane, respectively (Huang *et al.* 2008). The focal plane was set ~200 nm into the sample, so that depth ($z$) values of ~−200 nm corresponded to the substrate. Single-molecule images due to the transient entrance of individual BDP-TMR-alkyne molecules into the membrane phase (Sharonov and Hochstrasser 2006) were continuously recorded in the wide-field using an EM-CCD camera (iXon Ultra 897, Andor) at a fixed frame rate of 110 fps. A multifunction I/O board (PCI-6733, National Instruments) read the camera exposure timing signal, and accordingly modulated the RF synthesizer to achieve frame-synchronized stroboscopic excitation for the 560 nm laser in two modes. In the first mode, paired pulses of $\tau$ = ~1 ms duration and $\Delta t$ = 2.5 ms center-to-center separation were repeatedly applied across tandem camera frames (Figure 1g inset), so that single-molecule images from the paired odd-even frames captured transient displacements in the $\Delta t$ time window (Xiang *et al.* 2020). In the second mode, pulses of $\tau$ = 0.5-2 ms duration were applied at the middle of each frame (Figure 1h inset). Single-molecule displacements were thus assessed between consecutive frames for $\Delta t$ = 9.1 ms displacement time. The estimated peak and average power densities of the excitation laser at the sample were ~10 and ~1 kW/cm$^2$, respectively. 90,000-160,000 frames of single-molecule images were recorded under the above stroboscopic excitation schemes, from which $10^6$−$10^7$ molecules were accumulated across the view. The 488 nm laser was separately applied before and/or after the above stroboscopic single-molecule experiments for epifluorescence imaging of AF488-tagged CTB and GFP-tagged GRAMD1a and GRAMD2a.

**Concurrent SM*d*M and SR-SMLM.** Concurrent SM*d*M and SR-SMLM of Nile Red were achieved on another homebuilt system as described previously (Moon *et al.* 2017; Xiang *et al.* 2020). Briefly, frame-synchronized stroboscopic excitation was achieved through direct power modulation of the 561 nm laser (OBIS 561 LS, Coherent) using a multifunction I/O board (PCI-6733, National Instruments) (Xiang *et al.* 2020). Paired pulses of $\tau$ = 2 ms duration and $\Delta t$ = 2.5 ms center-to-center separation were repeatedly applied across tandem camera frames for evaluation of single-molecule displacements from the paired odd-even frames. The estimated peak and average power densities of the excitation lasers at the sample were ~6 and ~1 kW/cm$^2$, respectively. Single-molecule emission due to the transient entrance of individual Nile Red molecules into the membrane phase (Sharonov and Hochstrasser 2006) was split 50:50 for the concurrent recording of the unmodified images and the dispersed emission spectra in the wide-field(Moon *et al.* 2017). An EM-CCD camera (iXon Ultra 897, Andor) recorded ~120,000 frames at 110 fps.

**3D-SMLM of GRAMD1a and GRAMD2a.** For 3D-SMLM of GRAMD1a-AcGFP and GRAMD2a-AcGFP, transfected cells were fixed by 3% paraformaldehyde and 0.1% glutaraldehyde in phosphate-buffered saline (PBS) for 20 min, followed by a 5-min wash with freshly prepared 0.1% $NaBH_4$ in PBS. The sample was washed 3 times with PBS, and immunolabeled with a mouse anti-GFP primary antibody (Invitrogen A11120) and an Alexa Fluor 647-labeled secondary antibody (Invitrogen A21236). The labeled samples were imaged in a photoswitching buffer (100 mM Tris-HCl pH 7.5, 100 mM cysteamine, 5% glucose, 0.8 mg/mL glucose oxidase, and 40 µg/mL catalase) on the 3D-SMLM setup described above. ~50,000 frames of single-molecule images were recorded at 110 fps under continuous 647-nm excitation.

**Data analysis.** Single-molecule raw images were first localized and rendered into 3D-SMLM images as described previously (Rust *et al.* 2006; Huang *et al.* 2008). SR-SMLM data were processed as previously described (Moon *et al.* 2017; Zhang *et al.* 2015). SM*d*M analysis under the isotropic diffusion model was performed as described previously (Xiang *et al.* 2020), with the accumulated single-molecule displacements spatially binned into 100×100 nm$^2$ bins for MLE (maximum likelihood estimation) fits to a modified 2D random-walk model to obtain the local diffusion coefficient $D$. For the pSM*d*M analysis, for each spatial bin, we first calculated the angular coordinate $\varphi$ of each single-molecule displacement accumulated in the bin. As we consider molecules traveling in opposite directions to be diffusing along the same axis, we first flipped all displacements with $\varphi$ in the range of (-180°, 0°) by adding 180°, so that all the resultant $\varphi'$ values were in the range of [0°, 180°]. We then took the circular mean of $2\varphi'$ (in the range of [0°, 360°]) for all displacements in the spatial bin, and divided this value by 2 to obtain the average direction of diffusion $\theta$ (principal direction). Anisotropy $\alpha$ is also assessed in the process by dividing the modulus of the vector sum (in the $2\varphi'$ angle space) over the sum of the moduli of all the displacements, hence a value of 0 and 1 for fully isotropic and fully anisotropic (bidirectional along a line) diffusions, respectively. Results at each spatial bin were converted into a color for color-map presentation (*e.g.*, Figure 3g and Figure S1) *via* the HSV (hue, saturation, value) color space, with hue, saturation, and value corresponding to $\theta$, $\alpha$, and single-molecule count, respectively. The single-molecule displacements in each bin were next separately projected along and perpendicular to the calculated $\theta$ direction (*e.g.*, Figure 3bcef) for MLE fits to a modified 1D diffusion model (Xiang *et al.* 2020), with the larger resultant $D$ value of the two picked to generate pSM*d*M color maps.

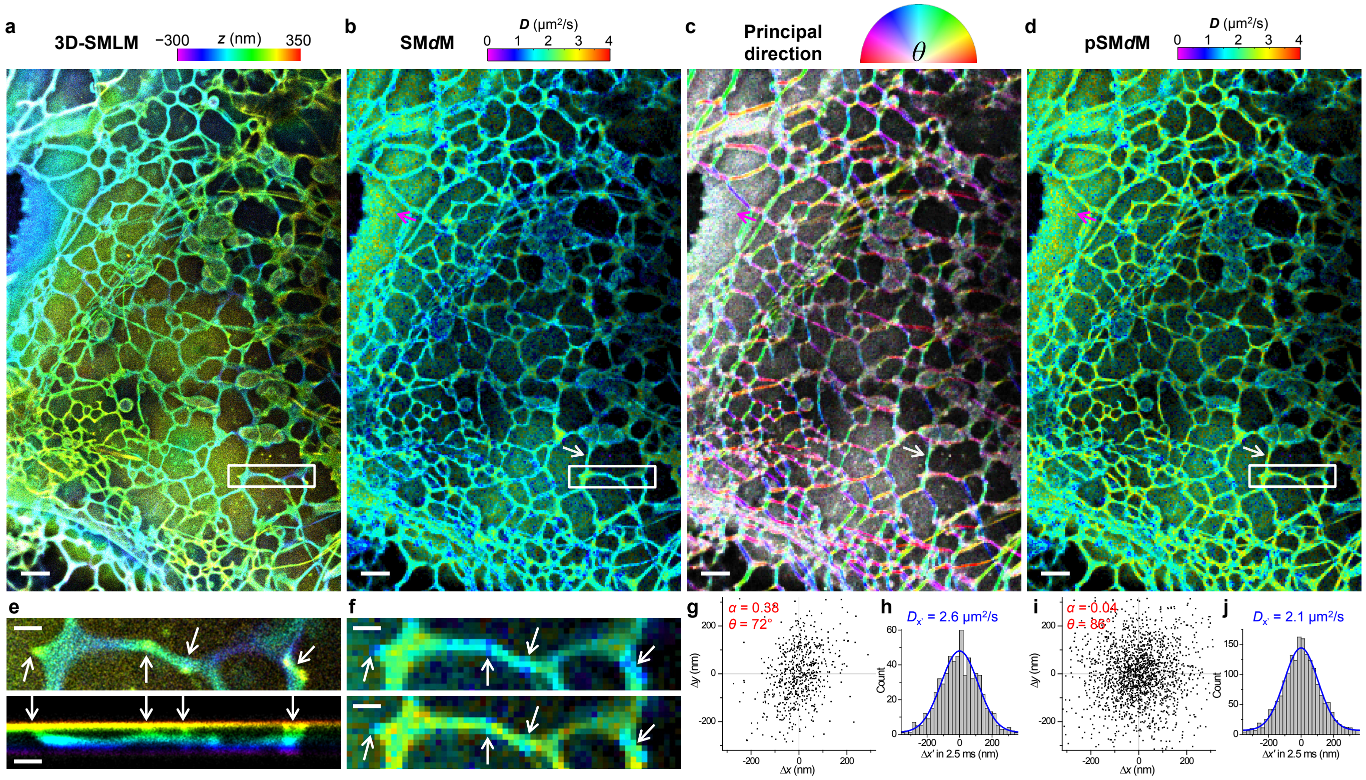


**Figure S1.** Additional stroboscopic PAINT and SM*d*M results of BDP-TMR-alkyne on wide-type COS-7 cell, acquired with tandem excitation pulses of $\tau$ = 1.5 ms duration and $\Delta t$ = 2.5 ms center-to-center separation. (a) 3D-SMLM image constructed from the single-molecule positions. Color presents depth (color scale on the top). (b) SM*d*M diffusivity map constructed from the single-molecule displacements of the same data, based on an isotropic 2D fitting model. Color presents $D$ value (color scale on the top). (c) Principal direction ($\theta$) - anisotropy ($\alpha$) color map of the diffusion data. Hue, saturation, and value present $\theta$, $\alpha$, and single-molecule count, respectively. (d) pSM*d*M diffusivity map after the principal-direction analysis. (e) Zoom-in of the white box in (a) (top), and its cross-sectional view in the *xz* direction (bottom). (f) Isotropic SM*d*M (top) and pSM*d*M (bottom) diffusivity maps of the same region. White arrows point to possible ER-PM contact sites. (g) 2D plot of single-molecule displacements in 2.5 ms for BDP-TMR-alkyne at an ER tubule [white arrows in (b-d)]. A principal direction $\theta$ of 72° is calculated with an anisotropy $\alpha$ of 0.38. (h) 1D distribution of the displacements in (g) projected along $\theta$. Blue curve: MLE fit to a 1D diffusion model with resultant $D$ = 2.6 μm$^2$/s. (i) 2D plot of single-molecule displacements in 2.5 ms for BDP-TMR-alkyne at the plasma membrane [magenta arrows in (b-d)]. $\alpha$ = 0.04 and $\theta$ = 86°. (j) 1D distribution of the displacements in (i) projected along $\theta$. Blue curve: MLE fit to a 1D diffusion model with resultant $D$ = 2.1 μm$^2$/s. Scale bars: 2 μm (a-d); 500 nm (e,f).

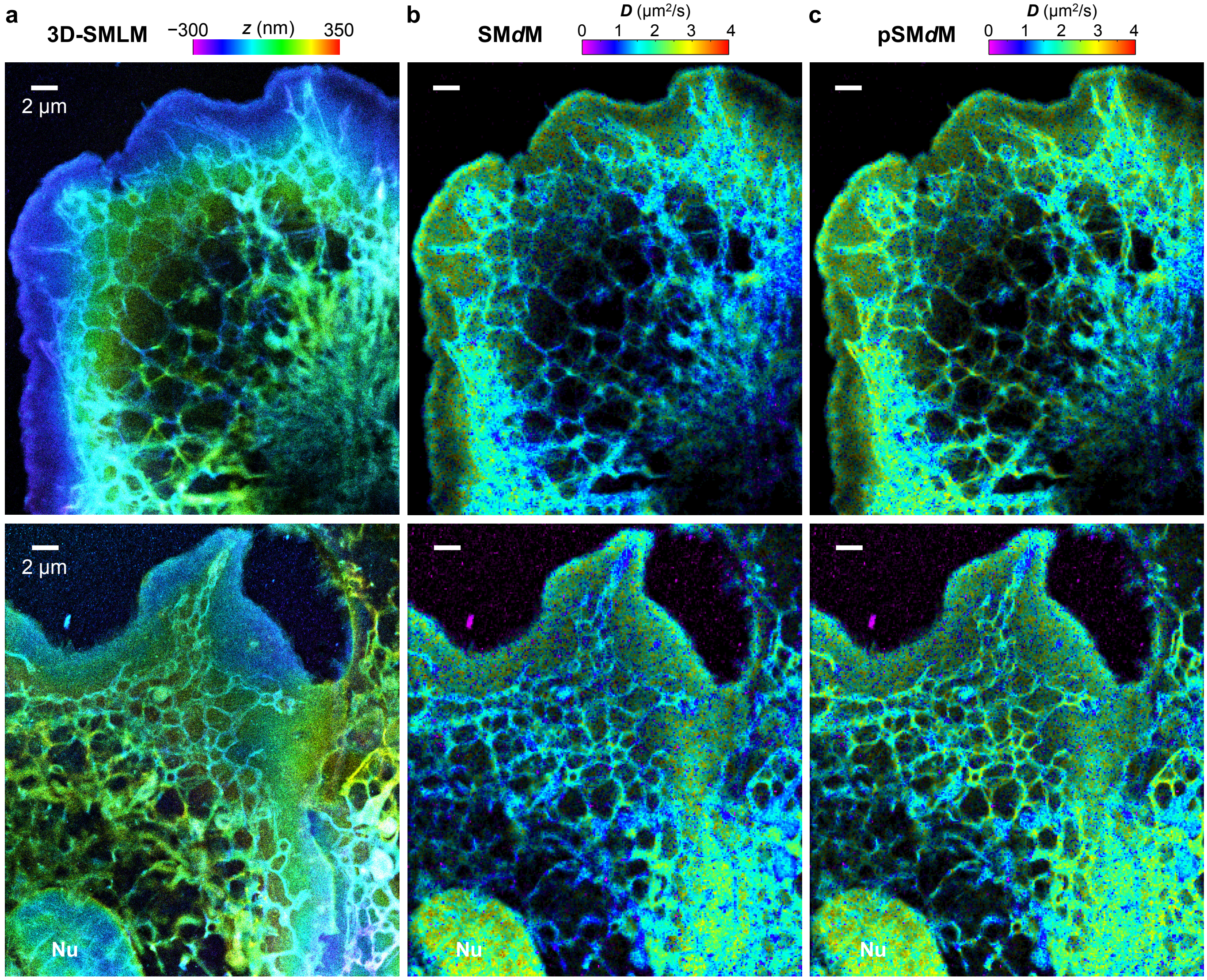


**Figure S2.** Additional stroboscopic PAINT and SM*d*M results of BDP-TMR-alkyne on wide-type COS-7 cell, acquired with stroboscopic excitations at the middle of every frame ($\Delta t$ = 9.1 ms), with excitation pulses of $\tau$ = 0.5 ms (top) and 2 ms (bottom) durations, respectively. (a) 3D-SMLM image. Color presents depth (color scale on the top). (b,c) Isotropic SM*d*M (b) and pSM*d*M (c) diffusivity maps. Color presents $D$ value (color scale on the top).
